# Assessing a causal relationship between circulating lipids and breast cancer risk: Mendelian randomization study

**DOI:** 10.1101/794594

**Authors:** Kelsey E. Johnson, Katherine M. Siewert, Derek Klarin, Scott M. Damrauer, the VA Million Veteran Program, Kyong-Mi Chang, Philip S. Tsao, Themistocles L. Assimes, Kara N. Maxwell, Benjamin F. Voight

## Abstract

**Objective:** To assess a potential causal relationship between genetic variants associated with plasma lipid traits (high-density lipoprotein cholesterol, HDL; low-density lipoprotein cholesterol, LDL; triglycerides, TG) with risk for breast cancer.

**Design:** Mendelian randomization (MR) study.

**Setting and Participants:** Data from genome-wide association studies in up to 215,551 subjects from the Million Veterans Project were used to construct genetic instruments for plasma lipid traits. The effect of these instruments on breast cancer risk was evaluated using genetic data from the BCAC consortium based on 122,977 breast cancer cases and 105,974 controls.

**Exposures:** Genetically modified plasma levels of LDL, HDL, or TG.

**Main Outcomes and Measures:** Odds ratio (OR) for breast cancer risk per standard-deviation increase in HDL, LDL, or TG.

**Results:** We observed that a 1-SD genetically determined increase in HDL levels is associated with an increased risk for all breast cancers (HDL: OR=1.08, 95% CI=1.04-1.13, P=7.4×10^−5^).

Multivariable MR analysis, which adjusted for the effects of LDL, TG, body mass index, and age at menarche, corroborated this observation for HDL (OR=1.06, 95% CI=1.03-1.10, P=4.9×10^−4^) and also a relationship between LDL and breast cancer risk (OR=1.03, 95% CI=1.01-1.07, P=0.02). We did not observe a difference in these relationships when stratified by breast tumor estrogen receptor status. We repeated this analysis using genetic variants independent of the leading association at core HDL pathway genes and found that these variants were also associated with risk for breast cancers (OR=1.11, 95% CI=1.06–1.16, P=1.5×10^−6^), including gene-specific associations at *ABCA1, APOE-APOC1-APOC4-APOC2* and *CETP*. In addition, we find evidence that genetic variation at the ABO locus affects both lipid levels and breast cancer.

**Conclusions:** Genetically elevated plasma HDL levels appear to increase breast cancer risk. Future studies are required to understand the mechanism underlying this putative causal relationship, with the goal to develop potential therapeutic strategies aimed at altering the HDL-mediated effect on breast cancer risk.

## Introduction

Breast cancer is the second leading cause of death for women, motivating the need for a better understanding of its etiology and more effective treatments.^1^ Cholesterol is a known risk factor for multiple diseases that have reported associations with breast cancer, including obesity, heart disease and diabetes.^2^ However, it is unknown whether cholesterol plays a causal role in breast cancer susceptibility.

The body of epidemiological and clinical trial studies to date has yet to clearly determine if there is a causal relationship between cholesterol and breast cancer. Observational epidemiological studies have reported positive, negative, or no relationship between lipid levels and breast cancer risk, however these studies can suffer from confounding.^3–5^ While recent evidence suggests that statin use may reduce breast cancer risk,^6^ cumulative meta-analyses are inconclusive.^7^ Recently, cholesterol-lowering medications have been associated with improved outcomes in breast cancer patients on hormonal therapy, suggesting an interaction of circulating cholesterol levels with estrogen-sensitive breast tissues.^8^ These mixed findings motivate the need for a high-powered causal inference analysis of lipids on breast cancer.

To try to resolve these discrepancies, recent studies have applied the framework of Mendelian randomization (MR) to determine if genetically elevated lipid levels associate with breast cancer risk. In a small sample of 1,187 breast cancer cases, Orho-Melander *et al*. used multivariable MR to find suggestive evidence of a relationship between triglycerides and breast cancer, but no association between LDL-cholesterol or HDL-cholesterol and cancer.^9^ In a second study, Nowak *et al*.^10^ performed an MR analysis with genetic association data from large GWAS for lipids and breast cancer.^11,12^ They reported nominal positive associations between LDL-cholesterol levels and all breast cancers, and between HDL-cholesterol levels and ER-positive breast cancers. While compelling, this study also had limitations. First, they used relatively few variants in their genetic instrument due to the need for addressing instrument heterogeneity and genetic correlation between lipid traits, resulting in a conservative analysis. Second, they analyzed each lipid trait separately rather than take advantage of multivariable methods to consider lipid traits together along with additional, potentially confounding causal risk factors. Third, the authors did not quantitatively assess heterogeneity to determine if the observed lipid associations were statistically different across breast cancer subtypes.

These studies motivate a Mendelian randomization study that considers multiple lipid traits concurrently to delineate the independent effect of each lipid trait on breast cancer susceptibility. Such an approach that includes the effects of all biomarkers plus confounding factors obviates the need to remove pleiotropic variants and the loss of statistical power which results from this removal. An ideal approach would consider the effects of body mass index (BMI) and age at menarche, known risk factors for breast cancer that are correlated with lipids.^13–19^

In what follows, we apply the causal inference framework of MR to determine if genetically elevated lipid traits modify breast cancer susceptibility, independent of one another and of BMI and age at menarche. We take advantage of a recent GWAS for lipid levels performed in up to 215,551 individuals of European ancestry^20^ to provide power for our causal inference analyses. Using a joint, multivariable MR, we find that genetic elevation of HDL trait increases risk for breast cancer, even after accounting for the effects of our genetic instruments on BMI and age at menarche. In addition, we perform a gene-specific Mendelian randomization, which focuses on variants in genes most central to the metabolism of cholesterol. This gene-specific approach supports HDL as a causal risk factor for breast cancer. Finally, we perform genetic correlation analyses to look for both genome-wide and locus-based correlation in effect sizes between lipids and breast cancer. Our results suggest that genetically elevated levels of HDL, and genetically lowered levels of TG, raise breast cancer risk.

## Methods

### Study Populations

Lipids GWAS summary statistics were obtained from the Million Veteran Program (MVP) (up to 215,551 European individuals)^20^ and the Global Lipids Genetics Consortium (GLGC) (up to 188,577 genotyped individuals).^12^ As additional exposures in multivariable MR analyses, we used BMI summary statistics from a meta-analysis of GWAS in up to 795,640 individuals, and age at menarche summary statistics from a meta-analysis of GWAS in up to 329,345 women of European ancestry.^16,21^ Genome-wide association study (GWAS) summary statistics from 122,977 breast cancer cases and 105,974 controls were obtained from the Breast Cancer Association Consortium (BCAC).^11^ More details on these cohorts are in the **Supplementary Methods**.

### Lipid meta-analysis

We performed a fixed-effects meta-analysis between each lipid trait (Total cholesterol (TC), LDL, HDL, and triglycerides) in GLGC and the corresponding lipid trait in the MVP cohort^12,20^ using the default settings in plink.^22^ These meta-analysis results appear well calibrated (**Supplementary Figure 1**).

### Mendelian randomization analyses

Mendelian randomization analyses were performed using the TwoSampleMR R package (https://github.com/MRCIEU/TwoSampleMR).^23^ For all analyses, we used the two-sample MR framework, which means that the lipid, BMI, age at menarche and breast cancer genetic associations were estimated in different cohorts. Unless otherwise noted, MR results reported in this manuscript used inverse-variance weighting assuming a random effects model. SNPs associated with each lipid trait were filtered for genome-wide significance (P < 5×10^−8^) from the MVP lipid study^12^, and then removed SNPs in linkage disequilibrium (r^2^ < 0.001 in UK10K consortium).^24^ Each of these independent, genome-wide significant SNPs was termed a genetic instrument. To reduce heterogeneity in our genetic instruments for single trait MR, we employed a pruning procedure (**Supplementary Methods**). Genetic instruments used in single trait MR are listed in **Supplementary Table 1**. For multivariable MR experiments, we generated genetic instruments by first filtering the genotyped variants for those present across all datasets. For each trait and data set combination (Yengo *et al*. for BMI; Day *et al*. for age at menarche, MVP and GLGC for HDL, LDL, and TG) we then filtered for genome-wide significance (P < 5×10^−8^), and for linkage disequilibrium (r^2^ < 0.001 in UK10K consortium).^24^ We then tested for instrument strength and validity^25^, and removed instruments driving heterogeneity (**Supplementary Methods**). Genetic instruments used in multivariable MR are listed in **Supplementary Table 2**. As the MR methods and tests we employed are highly correlated, we did not apply a multiple testing correction to the reported P-values.

### Core HDL and LDL pathway genetic instrument development

We defined sets of core genes for HDL or LDL that met the following criteria: (1) their protein products are known to play a key role in HDL or LDL biology (plus *HMGCR* and *NPC1L1*, two targets of LDL lowering drugs, in the LDL gene set), and (2) there were conditionally independent lipid trait-associated variants within 100kb upstream or downstream of the RefSeq coordinates for the gene (or locus, in the case of *APOE-APOC1-APOC4-APOC2* and *APOA4-APOC3-APOA1*).^20^ We then used the conditional HDL or LDL association statistics from Klarin et al. for those genes in gene-specific MR analyses.^20^ The loci included in each set and the genetic instruments used in each locus-specific MR are listed in **Supplementary Table 3**. We performed a separate fixed effects inverse-variance weighted MR with the conditionally independent genetic instruments at each gene, and performed fixed effects inverse variance weighted meta-analysis of the results across HDL or LDL genes using the R package meta.^26^

### Genetic correlation analyses

We performed cross-trait LD Score Regression using the LDSC toolkit with default parameters, with the BCAC association statistics for breast cancer and our meta-analysis of GLGC and MVP for lipid associations. We used the ρ-Hess software for our local genetic correlation analysis,^27^ using the UK10K reference panel, and the LD-independent loci published in Berisa *et al*. to partition the genome.^28^ We used a Bonferroni significance threshold based on the number of these independent loci (1,704 loci). There was minor cohort overlap between the GLGC and Breast Cancer GWAS due to the EPIC cohort.^10^ We included this overlap when performing ρ-Hess, using the cross-trait LD score intercept to estimate phenotypic correlation.

## Results

### Single trait Mendelian randomization (MR) in breast cancer

We first performed single trait MR analyses using summary statistics from MVP^20^ for each of four lipid traits (*i.e.*, total cholesterol - TC, high density lipoprotein cholesterol - HDL, low density lipoprotein cholesterol - LDL, and triglycerides - TG) as the intermediate biomarkers and risk for all breast cancers as the outcome (**Supplementary Figure 2**). We observed a significant relationship between genetically elevated HDL and breast cancer risk (OR=1.10 per standard deviation of lipid level increase, 95% CI=1.04-1.17, P = 2.1×10^−3^) and genetically decreased triglyceride levels and breast cancer risk (OR=0.93, 95% CI=0.88-0.99, P=0.015; **Supplementary Table 4**). Sensitivity analyses identified heterogeneity (**Methods, Supplementary Table 5**), but there was no evidence of bias from directional pleiotropy (**Methods, Supplementary Table 6**). To mitigate concerns of instrument heterogeneity, we removed genetic variants from our genetic instrument for each lipid trait that were responsible for instrument heterogeneity (**Supplementary Methods**), and again observed a relationship with HDL cholesterol (OR=1.08, 95% CI=1.04-1.13, P=7.4×10^−5^) and triglycerides (OR=0.94, 95% CI=0.90-0.98, P=2.6×10^−3^) (**Figure 1, Supplementary Figures 3 and 4, Supplementary Table 7**). As HDL and TG are inversely correlated^14,29^, the opposing relationship between these two lipid traits and breast cancer could be expected in single-trait analyses.

**Figure 1.**
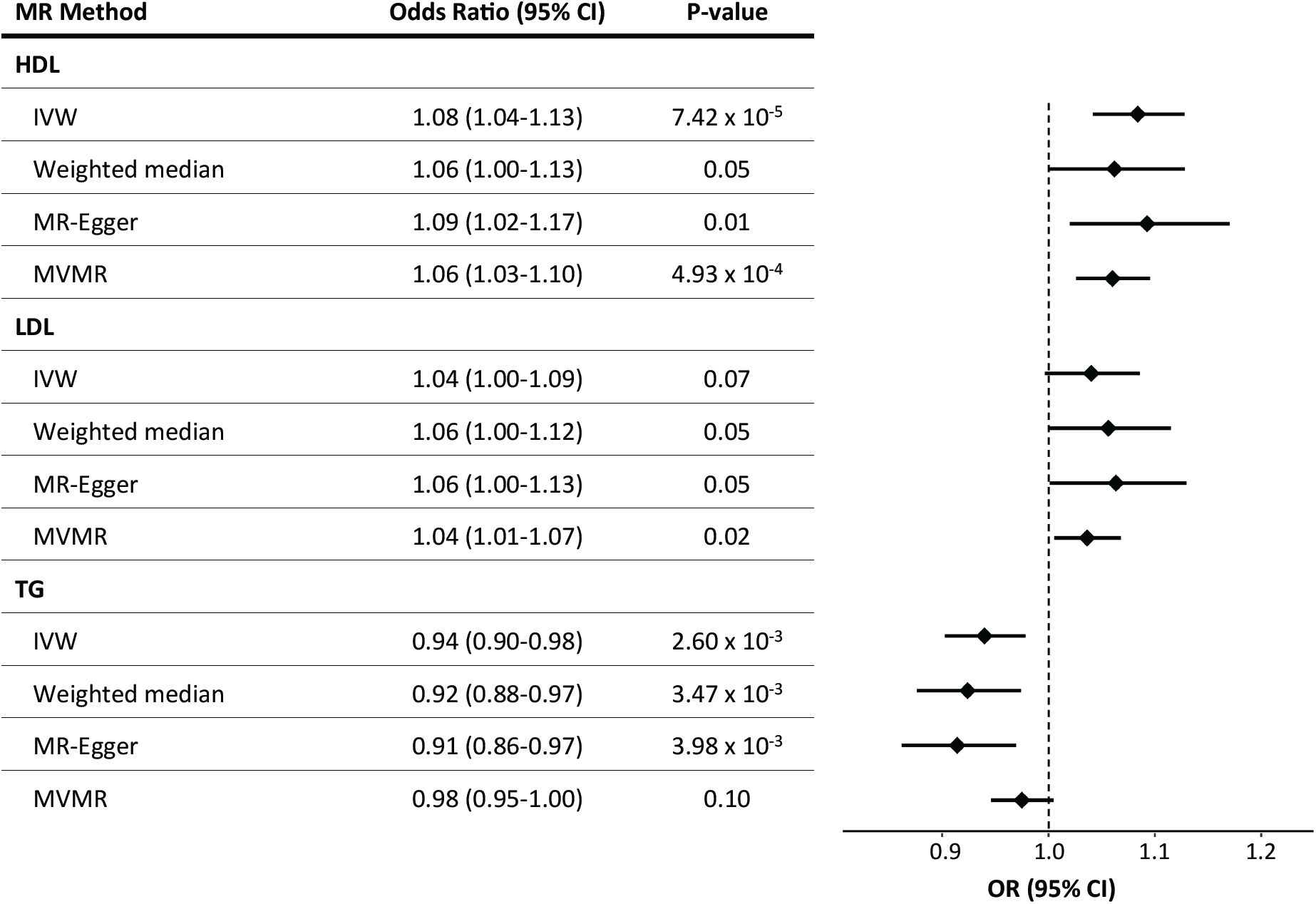
Results of MR analyses of the effects of HDL, LDL, or TG on breast cancer risk. IVW= inverse-variance weighted MR; MVMR = multivariable MR with HDL, LDL, TG, BMI, and age at menarche as exposures. Results plotted are after pruning for instrument heterogeneity.

We also tested the relationship between lipid traits and breast cancer using a meta-analysis of the two major lipid GWAS from MVP and GLGC, and GLGC alone. Overall, single trait MR analyses with the meta-analysis and GLGC lipid associations produced consistent results to those with MVP alone (**Supplementary Figure 5).** In a reciprocal single trait MR testing the effect of genetically-determined breast cancer risk on each lipid trait, we observed no relationship with HDL or LDL cholesterol (**Supplementary Table 8**), but did see a relationship with TG. However, a Steiger test for directionality confirmed that breast cancer as outcome was the correct causal direction for all lipid traits (**Supplementary Table 7**).^30^ We also performed genetic instrument pruning in the same manner as Nowak *et al*.: removing genetic instruments for LDL, HDL, and TG that were associated with at least one of the two other lipid traits (P<0.001).^10^ After this pruning, we did not find a significant relationship with LDL, HDL, or TG, and noted that this pruning procedure increased the size of the confidence intervals considerably (**Supplementary Table 9**).

### Multivariable Mendelian randomization with age at menarche and body mass index as exposures

It has been previously observed that body mass index (BMI) and age at menarche are both genetically correlated and epidemiologically associated with both breast cancer^19,31,32^ and lipid traits.^14,29^ To incorporate these potential confounders into our causal inference framework, we performed multivariable MR analyses using all three lipid traits (genetic effect estimates from MVP), age at menarche, and BMI as exposures, and breast cancer risk as the outcome (**Figure 1**). We observed relationships between genetically-influenced HDL, LDL, BMI, and age at menarche with breast cancer (HDL: OR=1.06, 95% CI=1.03-1.10, P=4.93×10^−4^; LDL: OR=1.04, 95% CI=1.01-1.07, P=0.02; BMI: OR=0.90, 95% CI=0.87-0.94, P=1.15×10^−6^; age at menarche: OR=0.96, 95% CI=0.93-0.99, P=2.44×10^−3^), but not TG (OR=0.98, 95% CI=0.95-1.00, P=0.10) (**Figure 1, Supplementary Table 10**). Our results were consistent before and after pruning for genetic instrument heterogeneity (**Supplementary Table 10)**, and when using summary statistics from three independent subsets of the breast cancer dataset (**Supplementary Figure 6, Supplementary Table 10**). We also performed multivariable MR with pairs of lipid traits with genetic effect estimates from different datasets (GLGC or MVP), with and without BMI, and saw consistent results (**Supplementary Table 11, Supplementary Figure 7**). Considering the genetic correlation between HDL and TG, the significant association of HDL compared to TG with breast cancer in multivariable analysis, and the consistent relationship between HDL and breast cancer across breast cancer datasets, we focused our further MR analyses on the relationship between HDL cholesterol and breast cancer, in addition to the previously reported association between LDL and breast cancer.^10^

### Mendelian randomization with outcome stratified by estrogen receptor status

We next performed a MR analysis of the relationship between genetically-influenced lipids and breast cancer risk stratified by estrogen receptor positive (ER+) or negative (ER-) status. We observed similar effect size estimates of the four lipid traits on the breast cancer subtypes as on breast cancer not stratified by subtype (**Supplementary Figure 8**). A formal test for heterogeneity found no evidence to reject the null hypothesis of homogeneity between the cancer subtypes (e.g. HDL: Cochran’s Q = 6.6×10^−5^, P = 0.99; **Supplementary Table 12**). Thus, we observed no substantive difference in the relationship from any lipid trait to ER+ or ER-breast cancers, consistent with the strong genetic correlation between these two breast cancer subtypes (cross-trait LD Score regression genetic correlation estimate = 0.62, P = 2.9 × 10^−83^).

### HDL and LDL pathway gene-specific Mendelian randomization

To further clarify causal association, we next examined associations for breast cancer risk at genetic variants near core HDL and LDL genes. We identified conditionally independent associations at core genes, which we define as genes or loci previously annotated with a core role in the metabolism of each lipid trait, or an established drug target (HDL: *ABCA1, APOA4-APOC3-APOA1, APOE-APOC1-APOC4-APOC2, CETP, LCAT, LIPC, LIPG, PLTP, SCARB1*; LDL: *APOB, HMGCR, LDLR, LPA, MYLIP, NPC1L1, PCSK9*) (**Methods, Supplementary Table 3**). For each gene or locus with at least two conditionally independent genetic instruments (all except *LCAT* and *MYLIP*), we performed inverse variance weighted MR (fixed effects model) with conditional HDL or LDL effect size estimates as the exposure and breast cancer risk as the outcome (**Supplementary Figures 9 and 10**). We observed a positive relationship between HDL and breast cancer risk at three loci (*ABCA1, APOE-APOC1-APOC4-APOC2, CETP*; **Figure 2**), and between LDL and breast cancer risk at one locus (*HMGCR*, **Supplementary Figure 11**). Combining the effect estimates across core genes in a meta-analysis, we observed a positive relationship for HDL (OR = 1.11, 95% CI = 1.06-1.16, P = 1.53×10^−6^; **Figure 2**) and LDL (OR = 1.07, 95% CI = 1.01-1.14, P = 0.02; **Supplementary Figure 11**). There was no evidence of heterogeneity across loci in either meta-analysis (HDL: Q=6.63, P=0.47; LDL: Q=5.53, P=0.35).

**Figure 2.**
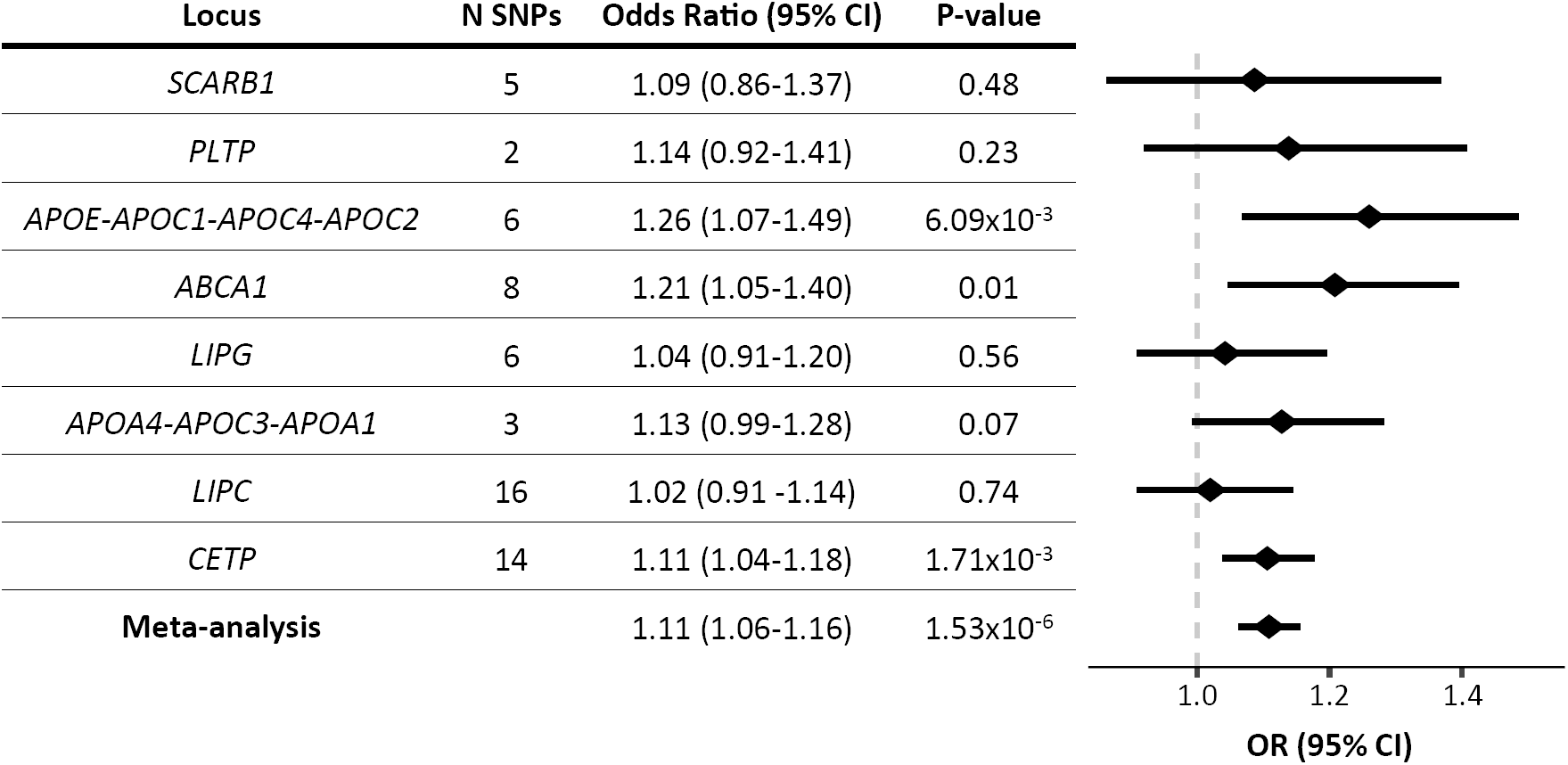
MR results for HDL gene-specific instruments (see **Supplementary Table 3**) and meta-analysis of effect estimates across genes.

### Genome-wide and local genetic correlation

If cholesterol levels were a causal risk factor for breast cancer, we might expect a correlation between the strength of genetic association with these two traits at genetic variants across the entire genome, in addition to those at genome-wide significant loci. To answer this question, we utilized two approaches to estimate genetic correlation between breast cancer and lipid traits. Consistent with our MR results, we found positive genetic correlation estimates for TC, LDL, and HDL; and a negative estimate for TG using cross-trait linkage disequilibrium score regression (**Supplementary Figure 12**).^14^ However, most P-values were not significant; the only nominal association (P < 0.05) was with total cholesterol and ER-negative breast cancer (P=0.04). In contrast, the ρ-Hess method, which uses a different model of genetic correlation, found significant (P<0.05) correlations between all four lipid traits and breast cancer, with directions consistent with our MR results (**Supplementary Table 13**).

To discover specific new loci that may be specifically correlated, we used the ρ-Hess method to search for genomic regions with genetic correlation between this same lipid meta-analysis and breast cancer.^27^ ρ-Hess identified one region that surpassed Bonferroni test correction, with a positive correlation between both LDL and TC and breast cancer (**Supplementary Table 14**). In this region there are two SNPs in high LD (rs532436 and rs635634, r^2^=0.99) that are genome-wide significantly associated with LDL (rs532436: P=9.98× 10^−112^), TC (P=1.68 × 10^−98^) and breast cancer (P=2.9 × 10^−8^). These SNPs lie within an intron of the *ABO* gene and are associated with change in gene expression of *ABO* in multiple tissues, suggesting ABO is a possible causal gene for these trait associations.^33^

## Discussion

Causal inference is one of the most challenging problems in biology and medicine, requiring a strong consensus of evidence from multiple sources. Excitingly, large-scale human genetics data provides the opportunity to bring an additional line of evidence to this important problem. To clarify how circulating levels of serum lipids levels might influence risk of breast cancer, we utilized the framework of Mendelian randomization. We provide evidence that genetically elevated HDL and LDL levels increase the risk for breast cancer in support of a causal hypothesis.

While substantial effort has been spent developing HDL raising therapies for cardiovascular disease prevention, independent studies have proposed an increase in all-cause mortality in individuals with high HDL levels.^34^ Our results suggest that therapies that aim to reduce cardiovascular risk by raising HDL levels might have an unintended consequence of elevated breast cancer risk. Specifically, our gene-based score using HDL-raising variation at the *CETP* locus predicted that *CETP*-based inhibition would elevate breast cancer risk (OR=1.11, 95% CI=1.04-1.18, P=1.71×10^−3^). Additionally, two recent Mendelian randomization studies reported causal evidence between elevated HDL and risk for age-related macular degeneration.^35,36^ These potential disease-increasing consequences may not have been possible to identify in safety trials, given the limited window of study to monitor progression or incidence of disease, the putative causal effect estimates, and the demographics of the study population (*i.e.*, higher proportion of male participants). Our result supports the use of human genetics data as both a novel strategy for therapeutic targeting and for the discovery of potential drug complications to direct long term post-clinical trial followup.^37^

Although Nowak *et al*. previously used Mendelian randomization to discover associations between lipids and breast cancer^9,10^, our report presents a reconsideration of these effects. Even after conditioning on the effects of HDL, BMI, and age at menarche, our MR analysis suggests a potential causal relationship between LDL and BC. Nowak *et al*. only found a relationship between HDL cholesterol and ER+ breast cancer, while we found a relationship between HDL cholesterol and risk for all breast cancers. We also find a previously unreported association with triglycerides and breast cancer, though our multivariable analysis suggests this may be explained by correlation between triglycerides and HDL, and not an independent triglyceride effect. In their analyses, Nowak *et al*. used a strict pruning procedure in an attempt to isolate the effects of each lipid trait. However, this approach reduces power due to the high genetic correlation of these traits. The multivariable approach taken here is an alternative way to estimate the effect of an exposure while accounting for correlated exposures.

We note several caveats to our analyses. The first is that MR makes a number of assumptions that must be met for accurate causal inference.^38,39^ Although we used statistical methods that try to detect and correct for violations of these assumptions, these methods are not guaranteed to correct for all types of confounding, and alternative causal inference frameworks outside of MR are warranted. We note that our effect estimates may be attenuated due to association of lipid IVS with the use of lipid-lowering medication. In addition, it is perhaps surprising that we did not find a significant genetic correlation between breast cancer and lipids using cross-trait LD score regression, however, our result corroborates a previous study which performed this analysis using smaller GWAS.^15^ Our lack of significant results could be caused by limited polygenicity of either trait, which decreases the power of this method.^14^ Lastly, we cannot be certain that the true underlying causal exposure is lipid levels, and not some phenotype for which lipids is a proxy. However, we are not aware of any process for which lipids is a proxy through which breast cancer would be affected.

The analyses presented here do not bring evidence on a specific mechanism for tumorigenesis, but they do bring renewed attention to potential mechanisms requiring future functional study. Cholesterol and its oxysterol metabolites, either in the circulatory system or in the local mammary microenvironment, may have direct effects on mammary tissue growth induction of breast tumorigenesis.^40,41^

Our study supports a causal relationship from increased HDL cholesterol to increased breast cancer risk, and this hypothesis warrants further exploration. Statins are widely prescribed to decrease LDL levels; however, statins also increase HDL levels. If further research substantiates the relationship between higher HDL levels and increased breast cancer risk, the consensus that HDL is “good cholesterol”, or of benign effect, may require re-evaluation.

## Acknowledgements

This work was supported by the US National Institutes of Health (R01 DK101478 to B.F.V., T32 GM008216 for K.E.J., T32 HG000046 for K.M.S.), and a Linda Pechenik Montague Investigator award (to B.F.V.). This research is based on data from the Million Veteran Program, Office of Research and Development, Veterans Health Administration and was supported by award #MVP000. This publication does not represent the views of the Department of Veterans Affairs or the United States Government. This research was also supported by two additional Department of Veterans Affairs awards (I01-BX003362 [Tsao/Chang], IK2-CX001780 [Damrauer]). This study makes use of data generated by the UK10K Consortium, derived from samples from the Avon Longitudinal Study of Parents and Children (ALSPAC) and the Department of Twin Research and Genetic Epidemiology (DTR). A full list of the investigators who contributed to the generation of the data is available from www.UK10K.org. Funding for UK10K was provided by the Wellcome Trust under award WT091310.

