## Supplementary information for "Assessing a causal relationship between circulating lipids and breast cancer risk: Mendelian randomization study"

KE Johnson, KM Siewert, *et al.*

**Supplementary Methods**

GWAS cohort details

Million Veteran Program (MVP) lipid summary statistics were estimated with European-ancestry individuals with the following sample sizes: 210,967 (HDL), 215,196 (LDL), 215,551 (total cholesterol), 211,491 (TG).[1] Lipids GWAS summary statistics from the Global Lipids Genetics Consortium (GLGC)[2] were downloaded from http://csg.sph.umich.edu/abecasis/public/lipids2013/ on March 3rd, 2017; this study included up to 188,577 genotyped individuals. Genome-wide association study (GWAS) summary statistics for breast cancer[3] from the Breast Cancer Association Consortium (BCAC) were downloaded from http://bcac.ccge.medschl.cam.ac.uk/bcacdata/oncoarray/gwas-icogs-and-oncoarray-summary-results/ on October 27th, 2017. This study performed a GWAS meta-analysis of a total of 122,977 breast cancer cases and 105,974 controls, and also breast cancer subtype meta-analyses with 69,501 cases (ER+) or 21,468 cases (ER-). In a multivariable MR experiment, we used summary statistics from three independent subsets of the BCAC consortium dataset: "Oncoarray", 61,282 female cases with breast cancer and 45,494 female controls; "iCOGS", 46,785 cases and 42,892 controls; and "GWAS", 14,910 cases and 17,588 controls from 11 GWAS. Breast cancer genome-wide association analyses were supported by the Government of Canada through Genome Canada and the Canadian Institutes of Health Research, the 'Ministère de l'Économie, de la Science et de l'Innovation du Québec' through Genome Québec and grant PSR-SIIRI-701, The National Institutes of Health (U19 CA148065, X01HG007492), Cancer Research UK (C1287/A10118, C1287/A16563, C1287/A10710) and The European Union (HEALTH-F2-2009-223175 and H2020 633784 and 634935). All studies and funders are listed in [3].

Heterogeneity analyses for single trait MR

To account for instrument heterogeneity in our raw lipid trait genetic instruments, we constructed a second genetic instrument subjected to the following pruning procedure to ensure the genetic instruments for each lipid trait met the assumption of no significant heterogeneity of effects.

Starting with a set of genetic instruments of all unlinked SNPs ($r^2 < 0.001$) reaching genome-wide significance ($P < 5 \times 10^{-8}$) for association with each lipid trait, we performed Cochran's Q test for heterogeneity[4], and removed SNPs with the largest contributions to Q until the null hypothesis of Cochran's Q test was no longer rejected ($P > 0.05$).

Multivariable MR tests of instrument strength and validity

We performed two heterogeneity tests[5], which are both modified Cochran's test of heterogeneity to evaluate instrumental variable strength ($Q_{X1}$) or validity ($Q_A$) in a two-sample (or more) summary data setting for multivariable MR. We found that our instrumental variables could predict each exposure trait when conditioning on the other exposure ($Q_{X1}$), but that there was significant heterogeneity suggesting invalid instruments when all significant genetic instruments were included ($Q_A$). To account for this, we further pruned our genetic instruments used for our multivariable analysis, using a stepwise post-hoc procedure to remove genetic instruments that contributed the most to $Q_A$ until the statistic was below the critical value ($\alpha = 0.05$).

**Supplementary Figures**

**Supplementary Figure 1**
QQ plot of Klarin et al and Willer et al meta-analysis results.

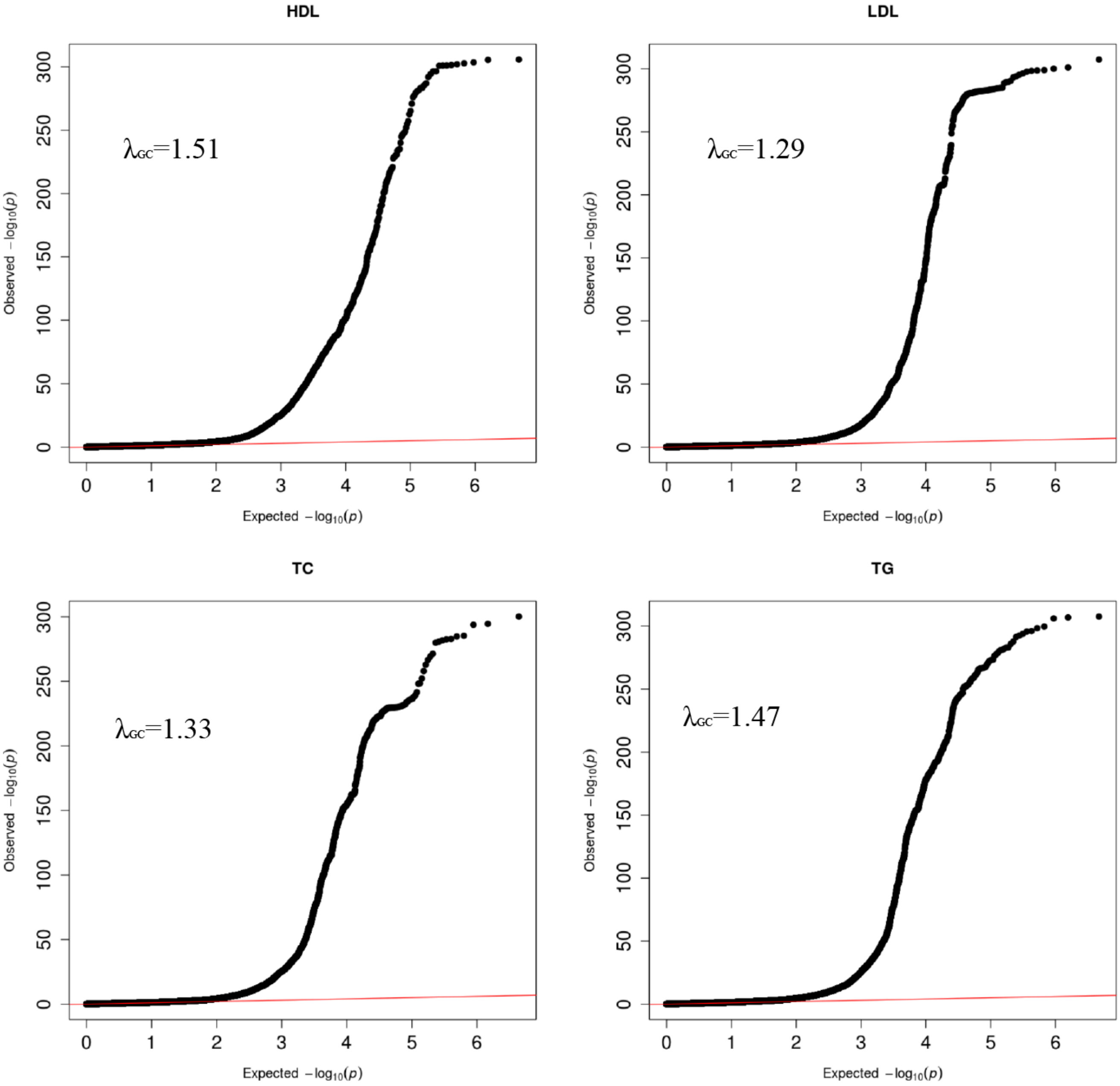

**Supplementary Figure 2**

Scatter plots of genetic instruments included in un-pruned single-trait MR analyses. Each plot contains effect estimates from MVP for one of four lipid traits (HDL, LDL, TC, TG) on the x-axis and effect estimates for risk of all breast cancers on the y-axis. Error bars represent the 95% confidence interval, and regression lines represent the slope estimate from one of three MR tests: inverse variance weighted (light blue), Egger regression (dark blue), weighted median (green).

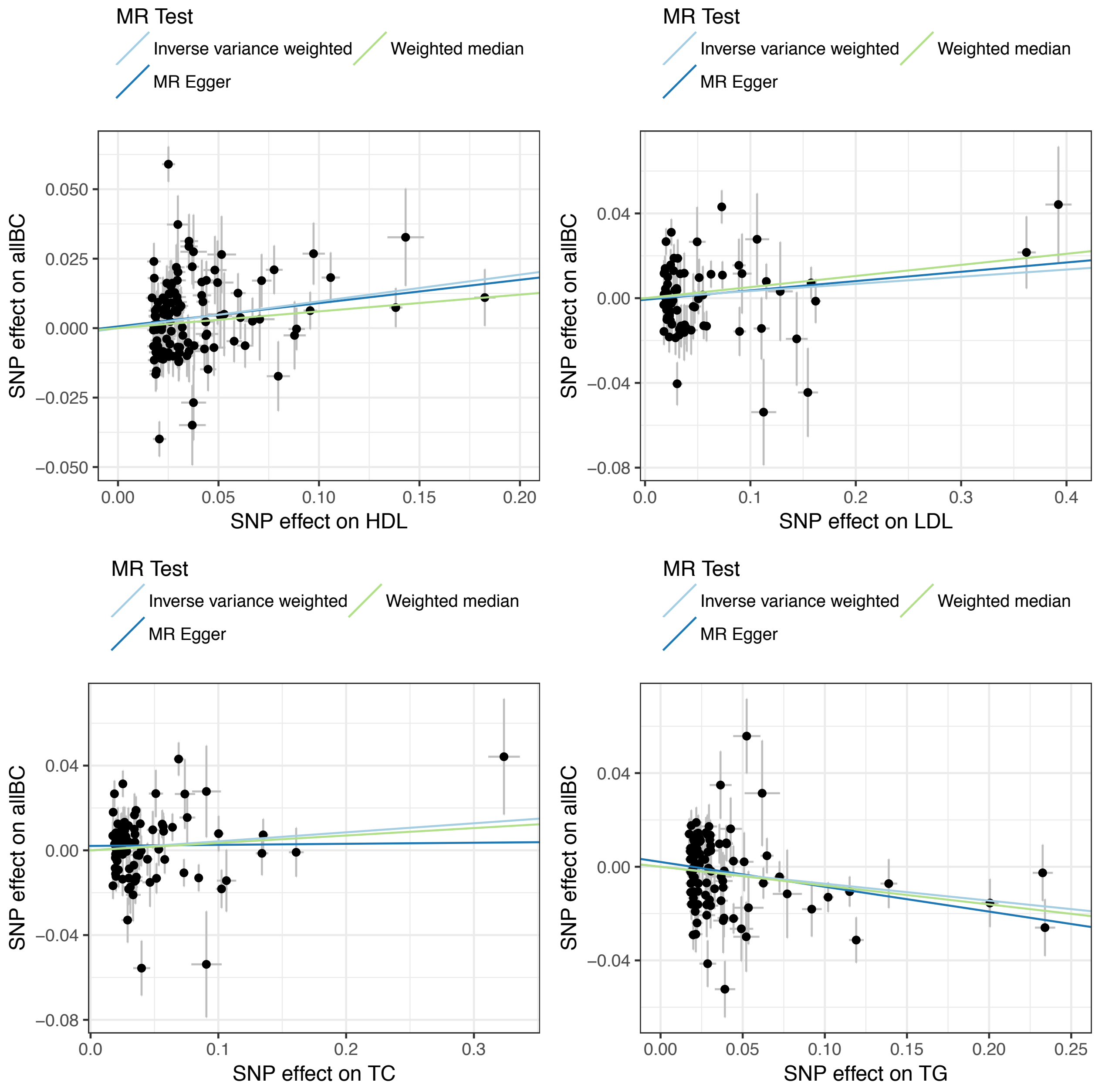

**Supplementary Figure 3**

Scatter plots of genetic instruments included in pruned single-trait MR analyses. Genetic instruments were pruned to pass heterogeneity test. Each plot contains effect estimates from MVP for one of four lipid traits (HDL, LDL, TC, TG) on the x-axis and effect estimates for risk of all breast cancers on the y-axis. Error bars represent the 95% confidence interval, and regression lines represent the slope estimate from one of three MR tests: inverse variance weighted (light blue), Egger regression (dark blue), weighted median (green).

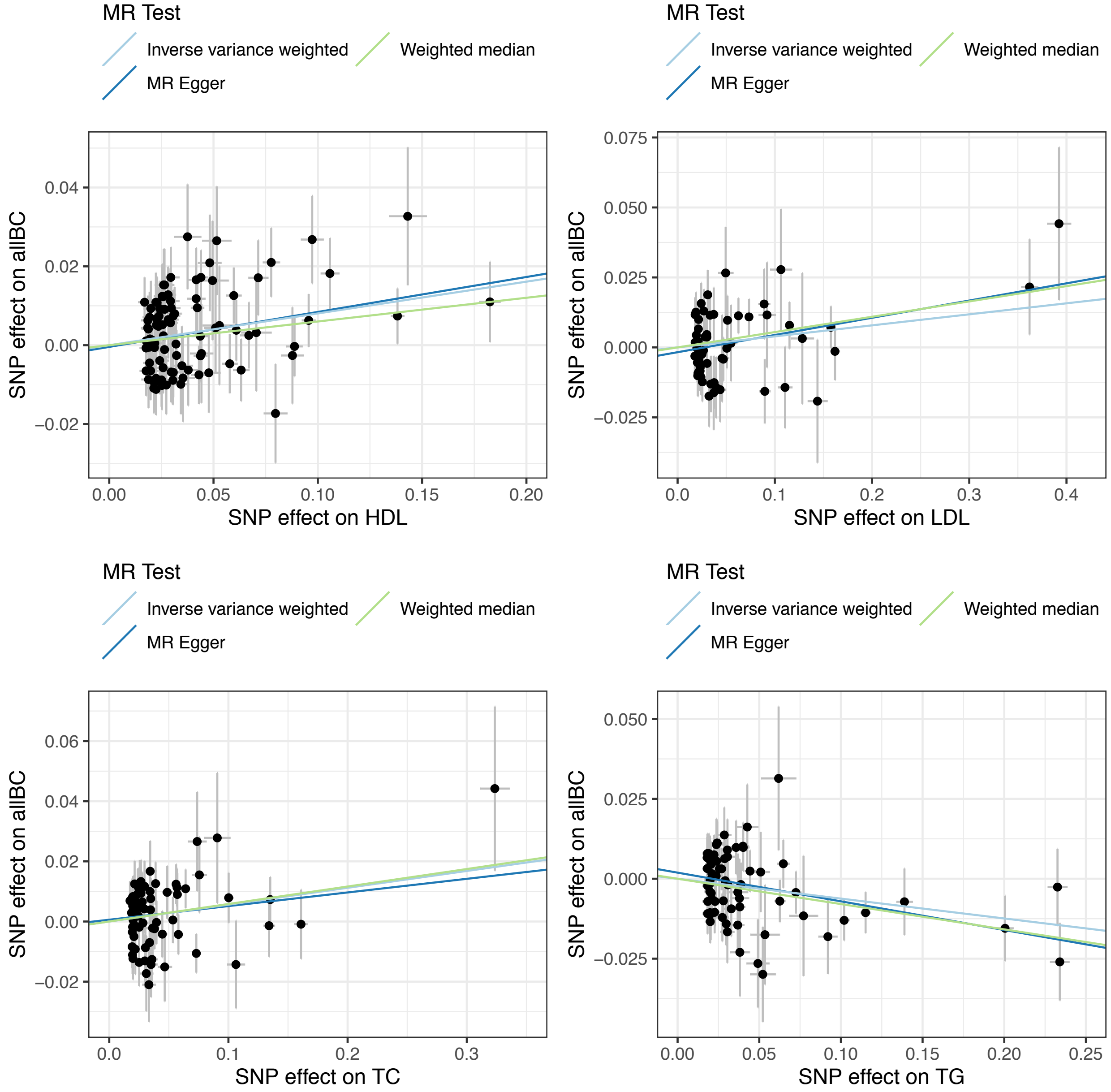

**Supplementary Figure 4**

Results of single trait MR analyses with a lipid trait as the exposure (data from MVP) and risk for all breast cancers as the outcome. Genetic instruments were pruned to pass heterogeneity test. Error bars represent the 95% confidence interval. Estimates were calculated using the inverse variance weighted (IVW) method. *P<0.05; **P<0.001.

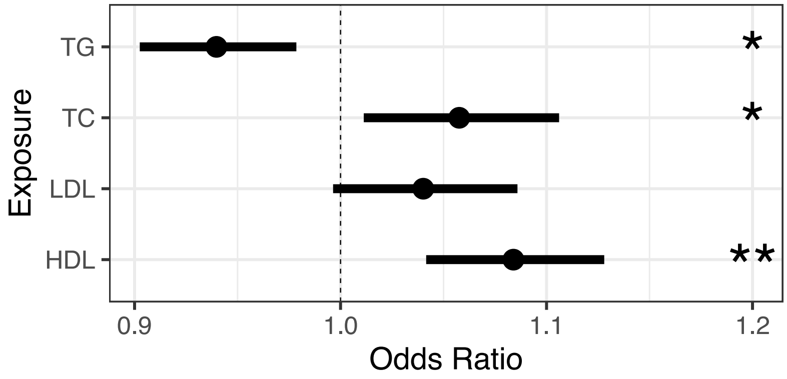

**Supplementary Figure 5**

Single-trait MR with lipid traits from three sources as the exposures (MVP, GLGC, and MVP+GLGC meta-analysis), and all breast cancers as the outcome. Genetic instruments were pruned to pass heterogeneity test. Error bars represent the 95% confidence interval. Estimates were calculated using the inverse variance weighted (IVW) method. *P<0.05; **P<0.001.

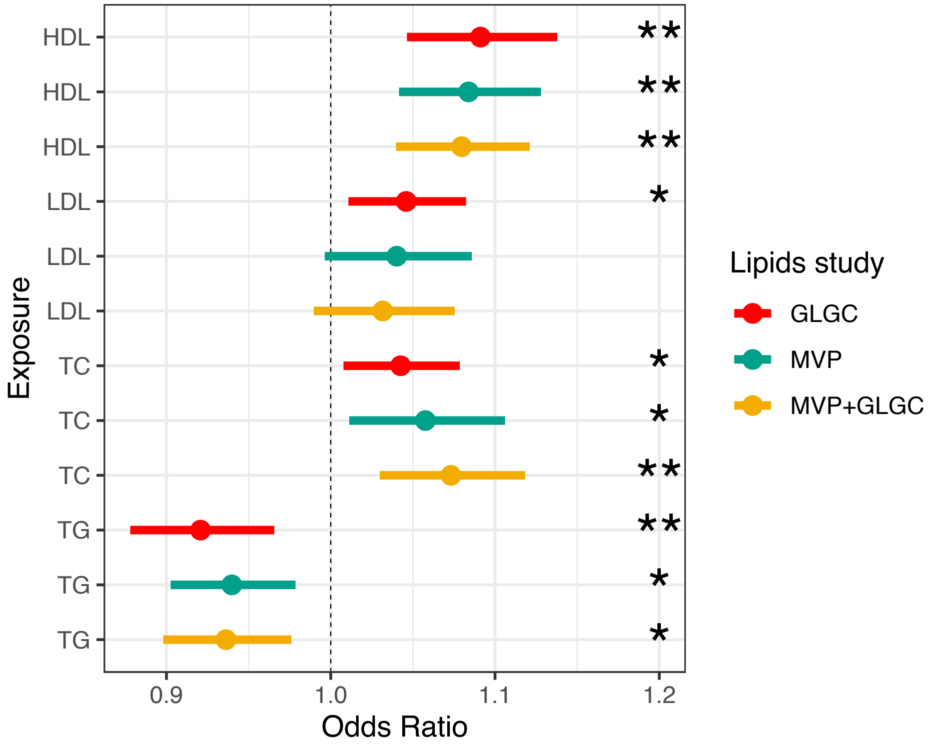

**Supplementary Figure 6**

Results of multivariable MR analyses with all three lipid traits, BMI, and age at menarche (AaM) as exposures, and breast cancer risk as the outcome. Each panel presents multivariable MR results using breast cancer summary statistics from an independent subset of the BCAC dataset (Oncoarray, iCOGS, or GWAS), or from the meta-analysis of all three together (BC meta-analysis) (**Supplementary Methods**). Results plotted are after pruning for instrument heterogeneity. Error bars represent the 95% confidence interval. Estimates were calculated using the inverse variance weighted (IVW) method. *P<0.05; **P<0.001.

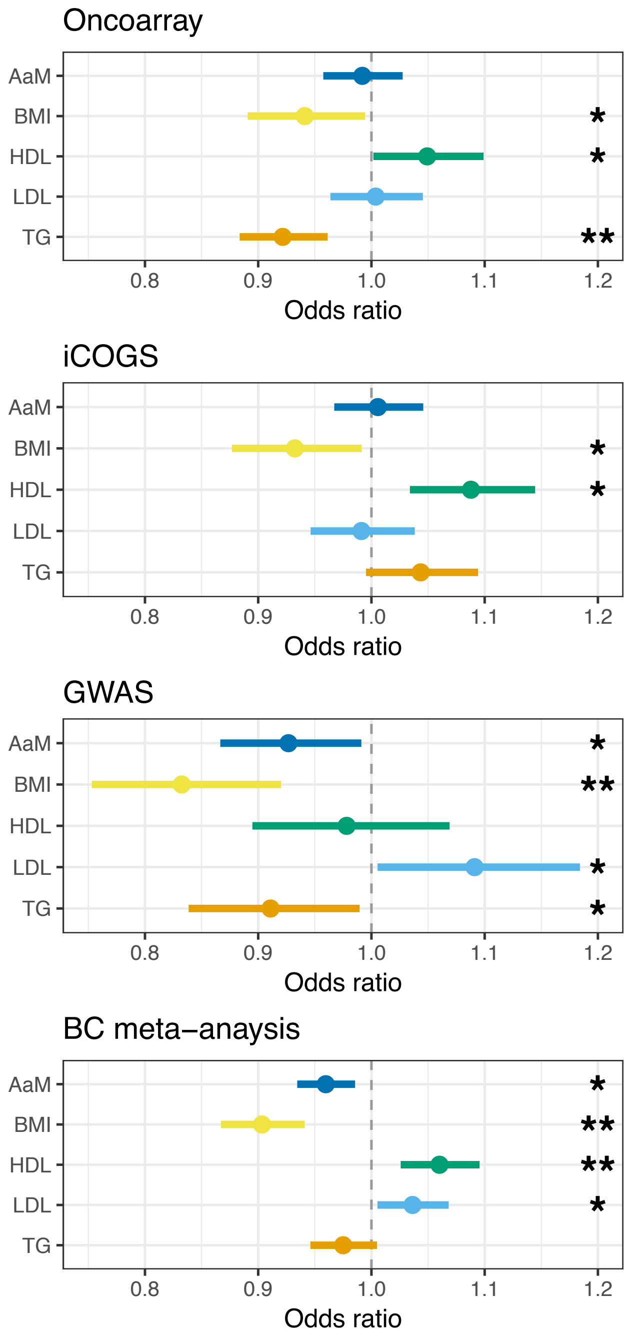

**Supplementary Figure 7**
Results of multivariable MR analyses including two lipid traits as exposures: **(A)** LDL and HDL, or **(B)** TG and HDL; with and without BMI as an additional exposure, and with risk for all breast cancers as the outcome. Results plotted are after pruning for instrument heterogeneity. The lipid effect estimates were from one of two GWAS datasets (MVP or GLGC), and the results of each combination of lipid datasets are in a single plot. Error bars represent the 95% confidence interval. Estimates were calculated using the inverse variance weighted (IVW) method. *P<0.05; **P<0.001.

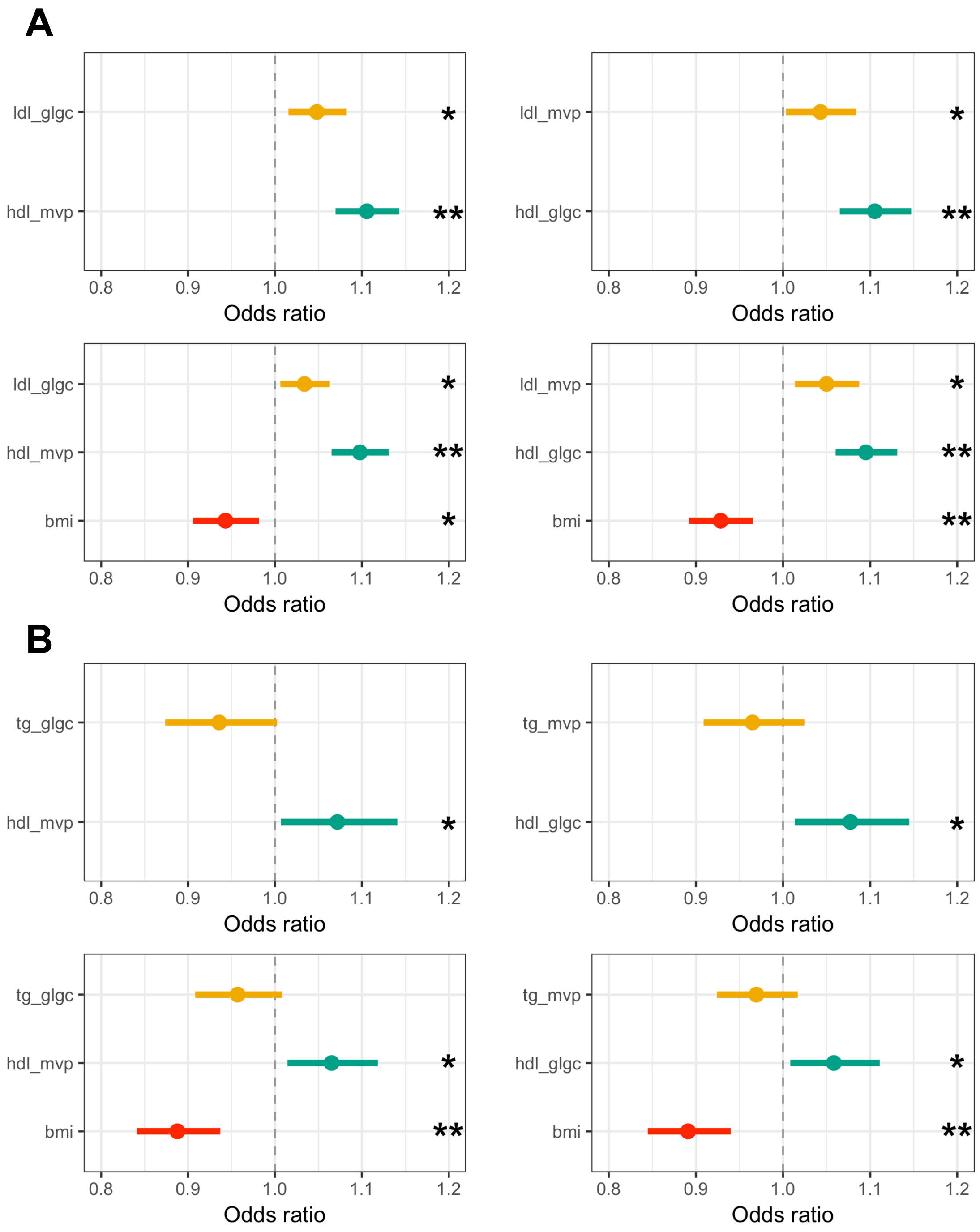

**Supplementary Figure 8**

Results of single trait MR with each lipid trait as an exposure, and one of three breast cancer traits as the outcome: all breast cancer, ER- breast cancers only, or ER+ breast cancers only. Error bars represent the 95% confidence interval. Estimates were calculated using a fixed-effects inverse variance weighted (IVW) method after pruning for instrument heterogeneity. Lipid association statistics come from the MVP data. **P<0.001, *P<0.05.

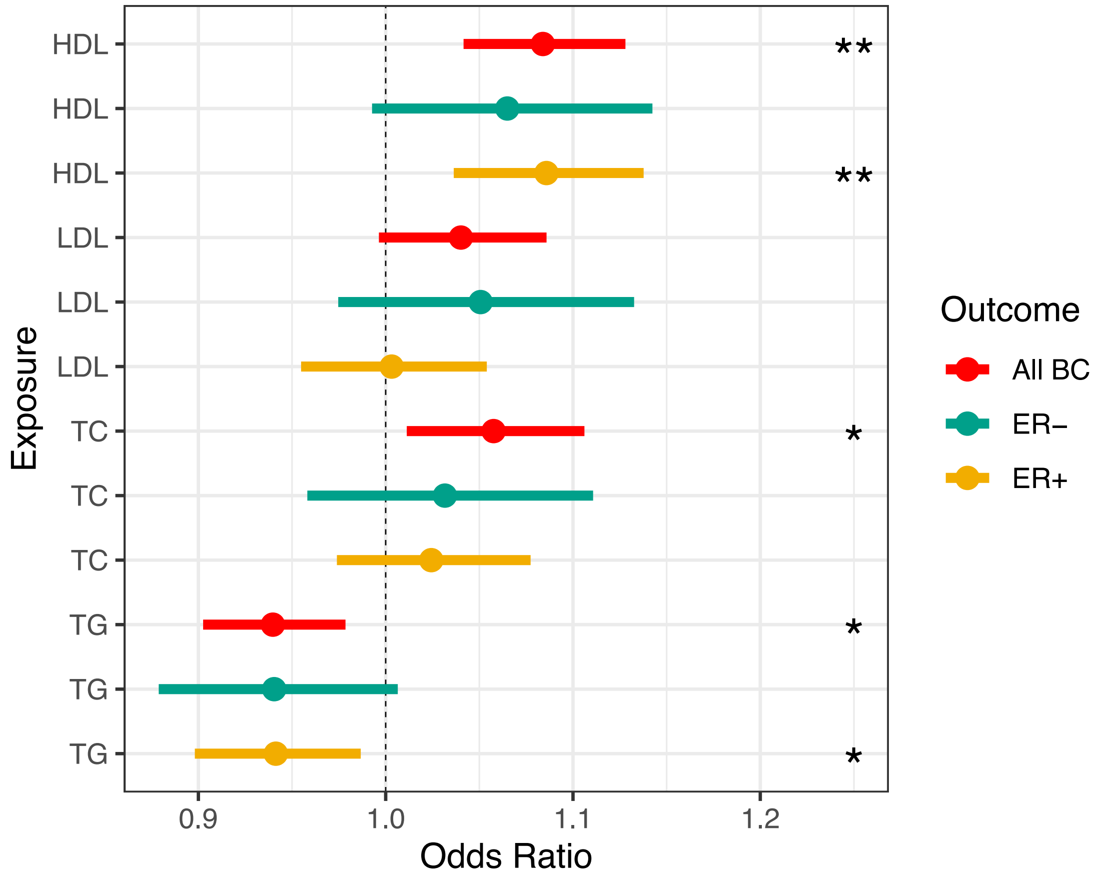

**Supplementary Figure 9**

Conditionally independent HDL-associated SNPs at canonical HDL metabolism pathway genes, plotted by their conditional effect estimates on HDL (from MVP) and effect estimates on all breast cancers. Error bars represent 95% confidence intervals. The dashed green line represents the regression line from fixed-effects IVW MR.

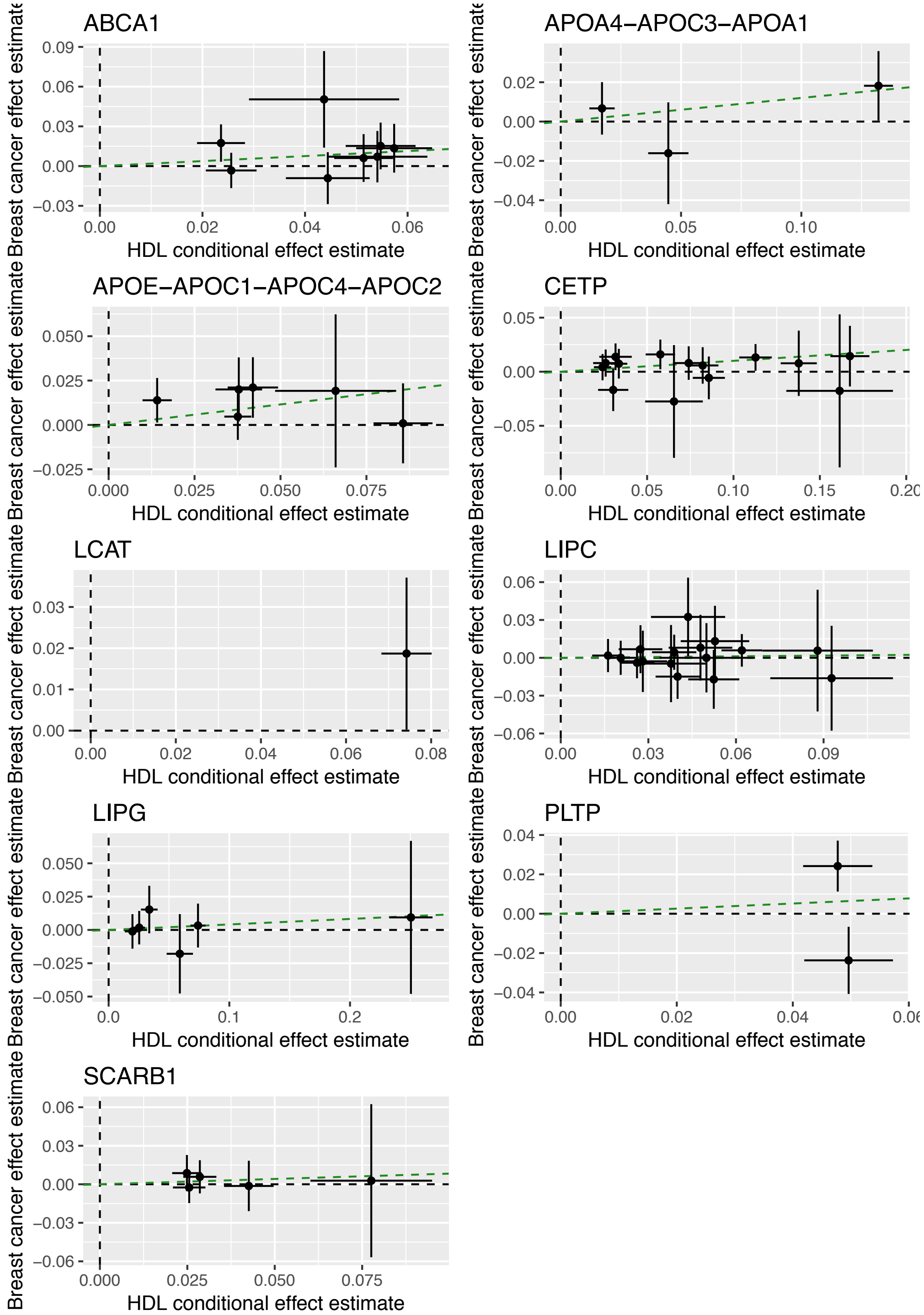

**Supplementary Figure 10**

Conditionally independent LDL-associated SNPs at canonical LDL metabolism pathway genes, plotted by their conditional effect estimates on LDL (from MVP) and effect estimates on all breast cancers. Error bars represent 95% confidence intervals. The dashed green line represents the regression line from fixed-effects IVW MR.

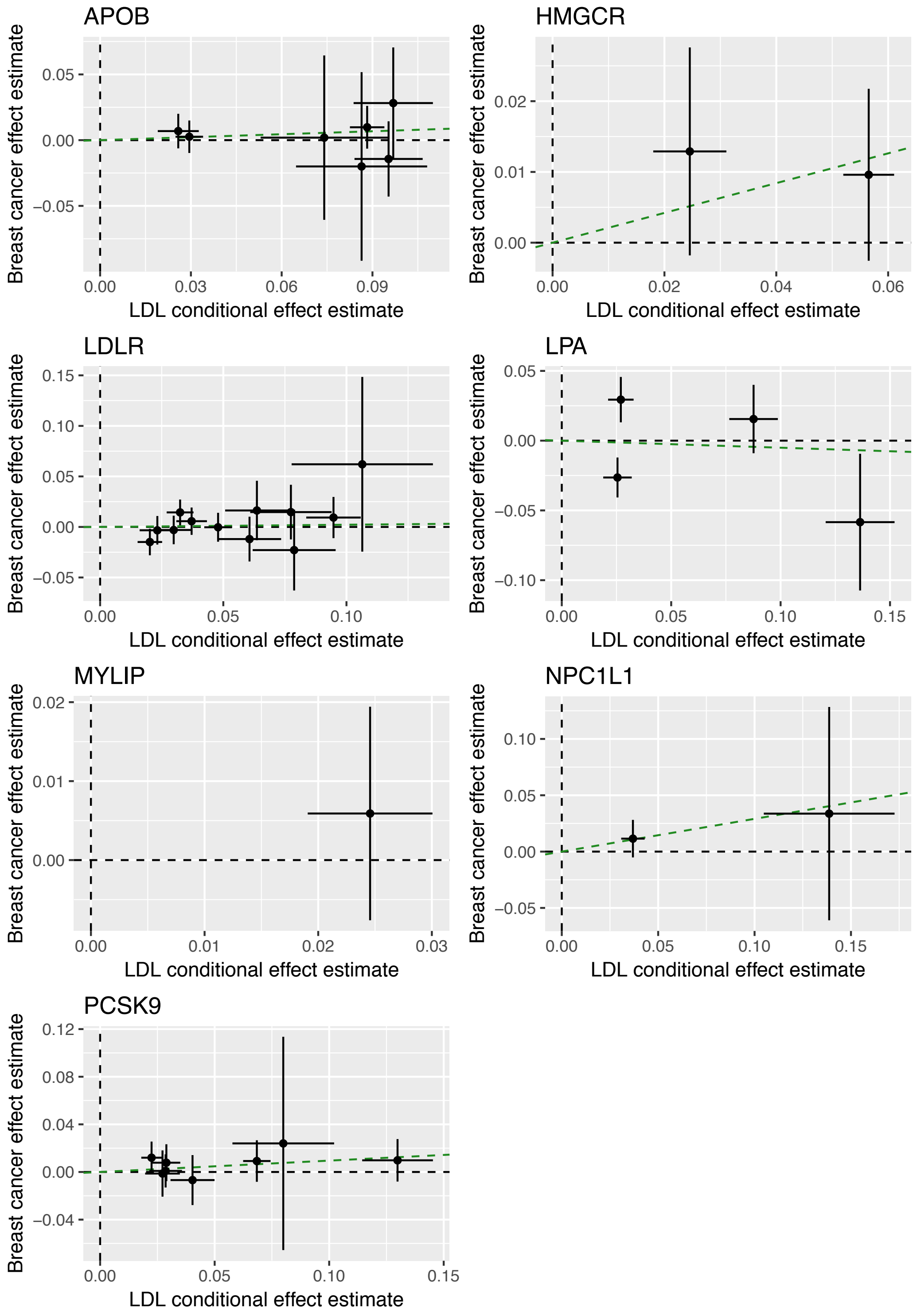

**Supplementary Figure 11**

Forest plot of MR results for LDL gene-specific instruments (see **Supplementary Table 4**) and meta-analysis of effect estimates across genes. Estimates were calculated using a fixed-effects inverse variance weighted (IVW) method.

| Gene | N SNPs | Odds Ratio (95% CI) | P-value |
|---|---|---|---|
| *NPC1L1* | 2 | 1.34 (0.92-1.95) | 0.13 |
| *HMGCR* | 2 | 1.23 (1.01-1.51) | 0.04 |
| *LPA* | 4 | 0.95 (0.78-1.15) | 0.61 |
| *APOB* | 7 | 1.08 (0.95-1.23) | 0.23 |
| *LDLR* | 11 | 1.02 (0.91-1.15) | 0.70 |
| *PCSK9* | 8 | 1.10 (0.99-1.23) | 0.08 |
| **Meta-analysis** | | 1.07 (1.01-1.14) | 0.02 |

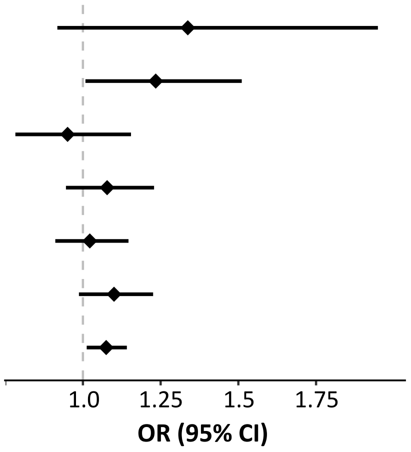

**Supplementary Figure 12**

Results of LD score regression testing for genetic correlation between each lipid trait and 3 breast cancer traits: all breast cancer, ER- breast cancers only, or ER+ breast cancers only. Error bars represent the 95% confidence interval. Lipid association statistics were from a meta-analysis of GLGC and MVP.

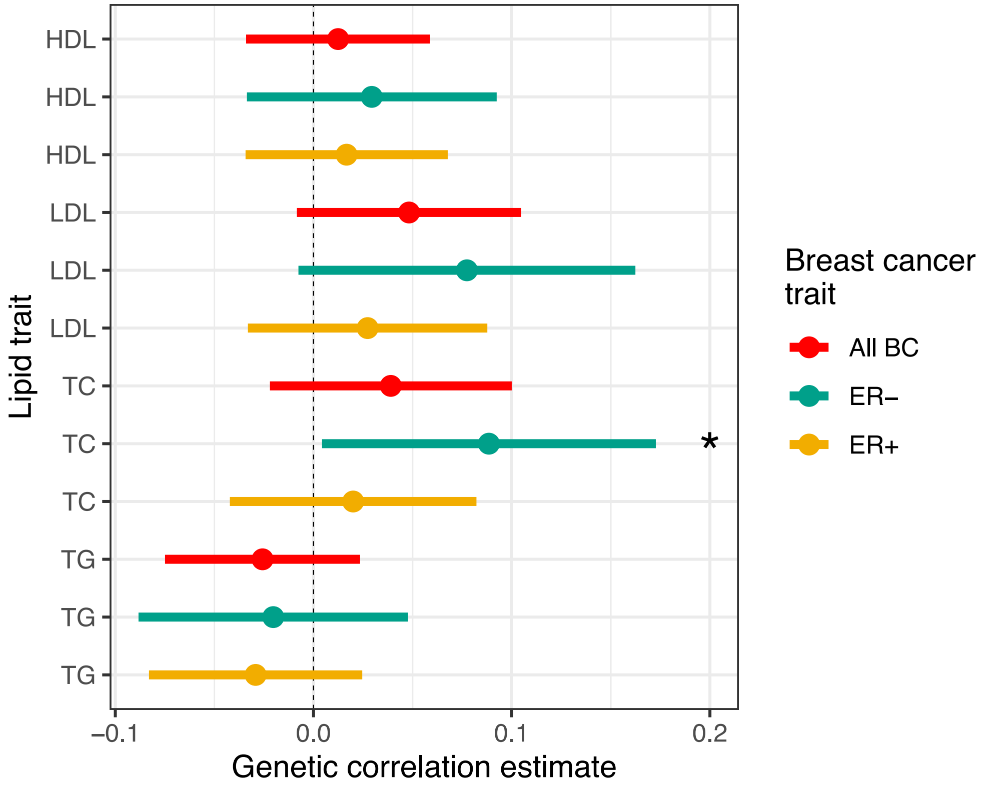

**Supplementary Table Descriptions**

**ST1.** Genetic instruments for single trait MR analyses. Lipid exposure summary statistics are from the MVP European dataset. inclPruned = was SNP included in pruned single trait MR analysis.

**ST2.** Genetic instruments used in multivariable MR analyses. expZ_* = summary statistics for trait listed as expZ (e.g. if exp1 is hdl_glgc, then exp1_beta is the effect size estimate for HDL from the GLGC dataset), bc_* = breast cancer summary statistics, test = unique identifier for each 2, 3, or 4-exposure MVMR experiment.

**ST3.** Conditionally independent variants used as genetic instruments in gene-specific MR for LDL or HDL. Lipid data are from a conditional analysis of summary statistics from MVP and GLGC meta-analysis[1].

**ST4.** Single trait MR results of with un-pruned lipid traits (MVP data) as the exposure and all breast cancers as the outcome, for a range of MR methods.

**ST5.** Heterogeneity analyses by Cochran's Q of un-pruned single-trait MR with MVP lipid data. Estimates are from the IVW method.

**ST6.** Pleiotropy analysis using Egger regression of un-pruned single-trait MR with MVP data.

**ST7.** Results of single trait MR, heterogeneity analyses, and directionality analyses with pruned lipid IV sets, using the IVW method with MVP data. Heterogeneity analyses use Cochran's Q. Directionality analyses use the Steiger test.

**ST8.** Results of a reciprocal single trait MR testing the effects of breast cancer as the exposure on each lipid trait as the outcome. Estimates are from the IVW method.

**ST9.** Results of single-trait MR analyses after pruning for genetic instruments associated with the other two listed lipid traits ($P<0.001$). Estimates are from the IVW method.

**ST10.** Results of multivariable MR with three lipid traits, BMI, and age at menarche (AaM) as exposures, and breast cancer risk as outcome. Results are from four separate multivariable MR experiments: three with summary statistics from independent subsets of the breast cancer dataset (Oncoarray, iCOGS, GWAS), and with the meta-analysis summary statistics (Meta). Lipid summary statistics are from MVP. Before/after pruning = results from MR before or after pruning for instrument heterogeneity.

**ST11.** Results of multivariable MR with 2 lipids, or 2 lipids and BMI, as exposures, and all breast cancers as outcome. Within a test, the effect estimates for each lipid trait are from different datasets (MVP or GLGC). Each test is separated by an empty row. Before/after pruning = results from MR before or after pruning for instrument heterogeneity.

**ST12.** Results of Cochran's Q test for heterogeneity in single trait IVW MR results for effect of a lipid trait on ER+ vs. ER- breast cancer.

**ST13.** Genome-wide correlation results using the ρ-Hess method.

**ST14.** Genomic regions with significant local genetic correlation between breast cancer and lipids using the ρ-Hess method. Only loci that passed Bonferroni correction ($P = 0.05/1703$ partitions $= 2.9 \times 10^{-5}$) are shown. Position = genomic coordinates of loci tested (hg19), N_SNPs = number of SNPs in partition, K = number eigenvectors used, Local-rhog= local genetic correlation.

**List of VA Million Veteran Program Contributors**

**MVP Executive Committee**

- Co-Chair: J. Michael Gaziano, M.D., M.P.H.
- Co-Chair: Rachel Ramoni, D.M.D., Sc.D.
- Jim Breeling, M.D. (ex-officio)
- Kyong-Mi Chang, M.D.
- Grant Huang, Ph.D.
- Sumitra Muralidhar, Ph.D.
- Christopher J. O'Donnell, M.D., M.P.H.
- Philip S. Tsao, Ph.D.

**MVP Program Office**

- Sumitra Muralidhar, Ph.D.
- Jennifer Moser, Ph.D.

**MVP Recruitment/Enrollment**

- Recruitment/Enrollment Director/Deputy Director, Boston – Stacey B. Whitbourne, Ph.D.; Jessica V. Brewer, M.P.H.
- MVP Coordinating Centers
    - Clinical Epidemiology Research Center (CERC), West Haven – John Concato, M.D., M.P.H.
    - Cooperative Studies Program Clinical Research Pharmacy Coordinating Center, Albuquerque - Stuart Warren, J.D., Pharm D.; Dean P. Argyres, M.S.
    - Genomics Coordinating Center, Palo Alto – Philip S. Tsao, Ph.D.
    - Massachusetts Veterans Epidemiology Research Information Center (MAVERIC), Boston - J. Michael Gaziano, M.D., M.P.H.
    - MVP Information Center, Canandaigua – Brady Stephens, M.S.
- Core Biorepository, Boston – Mary T. Brophy M.D., M.P.H.; Donald E. Humphries, Ph.D.
- MVP Informatics, Boston – Nhan Do, M.D.; Shahpoor Shayan
- Data Operations/Analytics, Boston – Xuan-Mai T. Nguyen, Ph.D.

**MVP Science**

- Genomics - Christopher J. O'Donnell, M.D., M.P.H.; Saiju Pyarajan Ph.D.; Philip S. Tsao, Ph.D.
- Phenomics - Kelly Cho, M.P.H, Ph.D.
- Data and Computational Sciences – Saiju Pyarajan, Ph.D.
- Statistical Genetics – Elizabeth Hauser, Ph.D.; Yan Sun, Ph.D.; Hongyu Zhao, Ph.D.

**MVP Local Site Investigators**

- Atlanta VA Medical Center (Peter Wilson)
  1670 Clairmont Rd, Decatur, GA 30033

- Bay Pines VA Healthcare System (Rachel McArdle)
  10,000 Bay Pines Blvd Bay Pines FL 33744
- Birmingham VA Medical Center (Louis Dellitalia)
  700 S. 19th Street Birmingham AL 35233
- Cincinnati VA Medical Center (John Harley)
  3200 Vine Street, Cincinnati, OH 45220
- Clement J. Zablocki VA Medical Center (Jeffrey Whittle)
  5000 West National Avenue, Milwaukee, WI 53295
- Durham VA Medical Center (Jean Beckham)
  508 Fulton Street Durham, NC 27705
- Edith Nourse Rogers Memorial Veterans Hospital (John Wells)
  200 Springs Road, Bedford, MA 01730
- Edward Hines, Jr. VA Medical Center (Salvador Gutierrez)
  5000 South 5th Avenue, Hines, IL 60141
- Fayetteville VA Medical Center (Gretchen Gibson)
  1100 N College Ave, Fayetteville, AR 72703
- VA Health Care Upstate New York (Laurence Kaminsky)
  113 Holland Avenue Albany NY 12208
- New Mexico VA Health Care System (Gerardo Villareal)
  1501 San Pedro Drive, S.E.Albuquerque, NM 87108
- VA Boston Healthcare System (Scott Kinlay)
  150 S. Huntington Avenue, Boston, MA 02130
- VA Western New York Healthcare System (Junzhe Xu)
  3495 Bailey Avenue Buffalo, NY 14215-1199
- Ralph H. Johnson VA Medical Center (Mark Hamner)
  109 Bee Street, Mental Health Research, Charleston, SC 29401
- Wm. Jennings Bryan Dorn VA Medical Center (Kathlyn Sue Haddock)
  6439 Garners Ferry Road, Columbia, SC 29209
- VA North Texas Health Care System (Sujata Bhushan)
  4500 S. LANCASTER ROAD, DALLAS, TX 75216
- Hampton VA Medical Center (Pran Iruvanti)
  100 Emancipation Drive, Hampton, VA 23667
- Hunter Holmes McGuire VA Medical Center (Michael Godschalk)
  1201 Broad Rock Blvd., Richmond, VA 23249
- Iowa City VA Health Care System (Zuhair Ballas)
  601 Highway 6 West, Iowa City, IA 52246-2208

- Jack C. Montgomery VA Medical Center (Malcolm Buford)
  1011 Honor Heights Dr., Muskogee, OK 74401
- James A. Haley Veterans' Hospital (Stephen Mastorides)
  13000 Bruce B. Downs Blvd., Tampa, FL 33612
- Louisville VA Medical Center (Jon Klein)
  800 Zorn Avenue, Louisville, KY 40206

- Manchester VA Medical Center (Nora Ratcliffe)
  718 Smyth Road, Manchester, NH 03104
- Miami VA Health Care System (Hermes Florez)
  1201 NW 16th Street, 11 GRC, Miami FL 33125
- Michael E. DeBakey VA Medical Center (Alan Swann)
  2002 Holcombe Blvd. Houston TX 77030
- Minneapolis VA Health Care System (Maureen Murdoch)
  One Veterans Drive Minneapolis MN 55417
- N. FL/S. GA Veterans Health System (Peruvemba Sriram)
  1601 SW Archer Road, Gainesville, FL 32608
- Northport VA Medical Center (Shing Shing Yeh)
  79 Middleville Road, Northport, NY 11768
- Overton Brooks VA Medical Center (Ronald Washburn)
  510 East Stoner Ave, Shreveport, LA 71101
- Philadelphia VA Medical Center (Darshana Jhala)
  3900 Woodland Avenue, Philadelphia, PA 19104
- Phoenix VA Health Care System (Samuel Aguayo)
  650 E. Indian School Road, Phoenix, AZ 85012
- Portland VA Medical Center (David Cohen)
  3710 SW U.S. Veterans Hospital Road, Portland, OR 97239
- Providence VA Medical Center (Satish Sharma)
  830 Chalkstone Avenue, Providence, RI 02908
- Richard Roudebush VA Medical Center (John Callaghan)
  1481 West 10th Street, Indianapolis, IN 46202
- Salem VA Medical Center (Kris Ann Oursler)
  1970 Roanoke Blvd.,Salem, VA 24153
- San Francisco VA Health Care System (Mary Whooley)
  4150 Clement Street, San Francisco, CA 94121
- South Texas Veterans Health Care System (Sunil Ahuja)
  7400 Merton Minter Boulevard, San Antonio, TX 78229
- Southeast Louisiana Veterans Health Care System (Amparo Gutierrez)
  2400 Canal Street, New Orleans, LA 70119
- Southern Arizona VA Health Care System (Ronald Schifman)
  3601 S 6th Ave, Tucson, AZ 85723
- Sioux Falls VA Health Care System (Jennifer Greco)
  2501 W 22nd St, Sioux Falls, SD 57105
- St. Louis VA Health Care System (Michael Rauchman)
  915 North Grand Blvd., St. Louis, MO 63106
- Syracuse VA Medical Center (Richard Servatius)
  800 Irving Avenue, Syracuse, NY 13210
- VA Eastern Kansas Health Care System (Mary Oehlert)
  4101 S 4th Street Trafficway, Leavenworth, KS 66048
- VA Greater Los Angeles Health Care System (Agnes Wallbom)

11301 Wilshire Blvd Los Angeles, CA 90073

- VA Loma Linda Healthcare System (Ronald Fernando)
  11201 Benton Street, Loma Linda, CA 92357
- VA Long Beach Healthcare System (Timothy Morgan)
  5901 East 7th Street Long Beach CA 90822
- VA Maine Healthcare System (Todd Stapley)
  1 VA Center, Augusta, ME 04330
- VA New York Harbor Healthcare System (Scott Sherman)
  423 East 23rd Street New York, NY 10010
- VA Pacific Islands Health Care System (Gwenevere Anderson)
  459 Patterson Rd, Honolulu, HI 96819
- VA Palo Alto Health Care System (Philip Tsao)
  3801 Miranda Avenue Palo Alto, CA 94304-1290
- VA Pittsburgh Health Care System (Elif Sonel)
  University Drive, Pittsburgh, PA 15240
- VA Puget Sound Health Care System (Edward Boyko)
  1660 S. Columbian Way Seattle, WA 98108-1597
- VA Salt Lake City Health Care System (Laurence Meyer)
  500 Foothill Drive Salt Lake City, UT 84148
- VA San Diego Healthcare System (Samir Gupta)
  3350 La Jolla Village Drive, San Diego, CA 92161
- VA Southern Nevada Healthcare System (Joseph Fayad)
  6900 North Pecos Road, North Las Vegas, NV 89086
- VA Tennessee Valley Healthcare System (Adriana Hung)
  1310 24th Ave. South Nashville, TN 37212
- Washington DC VA Medical Center (Jack Lichy)
  50 Irving St, Washington, D. C. 20422
- W.G. (Bill) Hefner VA Medical Center (Robin Hurley)
  1601 Brenner Ave, Salisbury, NC 28144
- White River Junction VA Medical Center (Brooks Robey)
  163 Veterans Drive, White River Junction, VT 05009
- William S. Middleton Memorial Veterans Hospital (Robert Striker)
- 2500 Overlook Terrace, Madison, WI 53705
